# The long-term application of composted organic wastes caused a lasting increase in vineyard soil health and multifunctionality

**DOI:** 10.1101/2025.10.22.683841

**Authors:** Lur Epelde, Ulas Karaoz, Carlos Garbisu, José Félix Cibriáin, Eoin Brodie

## Abstract

As a consequence of climate change and unsustainable management practices, agricultural soils in the Mediterranean region are often degraded. The application of composted organic wastes is traditionally considered a beneficial practice to improve soil health. We studied the permanence of the beneficial effects of the long-term (20 years) application of composted organic wastes, versus mineral fertilization or no fertilization, on Mediterranean vineyard soil health. To this aim, four years after ceasing fertilization, a comprehensive analysis of soil physicochemical and biological properties, including prokaryotic diversity and functional traits by genome-centric metagenomics, was conducted to ascertain whether the beneficial effects of organic fertilization were maintained 4 years after its end. In general, but not always, soils treated with composted organic wastes showed significantly higher values of many physicochemical and biological properties (e.g., organic matter, Olsen phosphorus and extractable potassium), resulting in improved soil multifunctionality. However, despite such statistical significance, the quantitative magnitude of many of those differences was small. Fertilization regimes had a lasting strong influence on soil prokaryotic communities, as 70% of the 200 metagenome-assembled genomes and 86% of the prokaryotic functional traits showed significant differences among treatments. Organic amendments promoted nitrifying taxa such as *Nitrosocosmicus oleophilus* and *Nitrospira japonica*. In minerally-fertilized soils, resource acquisition and stress tolerance strategies were fostered among prokaryotic communities, possibly due to resource limitation and soil degradation, respectively. Stress tolerance traits were lower under organic fertilization, likely due to the improved soil properties and functions. The results obtained suggest that soil functions are influenced by microbial genes involved in nitrogen and carbon cycling, underscoring the central role of microbial metabolism in sustaining soil health and ecosystem functioning. This study demonstrates the lasting benefits of composted organic amendments in promoting soil multifunctionality in vineyard soils.

## Introduction

Agricultural productivity and sustainability in the Mediterranean region are threatened by, among other causes, climate change, soil degradation, loss of biodiversity, and water resource depletion. These issues are exacerbated by poor management practices, including intensive tillage and the overuse of fertilizers and pesticides. Intensive agriculture typically contributes to the loss of soil organic matter (OM) and biodiversity, while promoting soil erosion, compaction, nutrient imbalance, and contamination.

In Mediterranean viticulture, soil health is often negatively impacted by, among other practices, frequent ploughing and an excessive use of phytosanitary products (Aranda et al., 2011; Calleja-Cervantes et al., 2015). Then, in the last years and decades, a soil health-based, more sustainable viticulture has been promoted by means of encouraging a variety of practices (e.g., the application of organic amendments to vineyard soil) aimed at protecting the integrity and resilience of the soil ecosystem, as well as to maximize the provision of soil ecosystem functions and services. In particular, the application of organic fertilizers can stimulate the activity of the soil biota, increase soil OM content and water-holding capacity, and enhance carbon storage and sequestration (Sharma et al., 2021). Still, organic fertilizers do not invariably enhance soil microbial diversity, and greater biodiversity does not necessarily indicate healthier soil.

Soil microbial communities are known to play a pivotal role in soil ecosystem processes and services, including nutrient cycling, carbon sequestration, and water regulation, while providing ecological stability (i.e., resistance and resilience) against biotic and abiotic disturbances, a crucial aspect in the current scenario of climate change and biodiversity loss (Barrios, 2007). Agricultural practices that enhance soil biodiversity, and specifically soil microbial diversity, are key for the functional sustainability of agricultural soils and systems (Wall et al., 2015). In particular, soil microorganisms are integral to a productive and functional viticulture and winemaking, influencing, among other aspects, grape quality and wine terroir (Griggs et al., 2021). Despite the crucial importance of soil microorganisms for a sustainable agriculture, the drivers of microbial diversity in vineyard soils still remain insufficiently understood and controversial (Hendgen et al., 2018). While organic fertilization commonly stimulates the biomass and activity of soil microorganisms, its effects on soil microbial diversity can vary widely depending on edaphoclimatic conditions, the composition of the organic products used as fertilizers, and the specific taxonomic groups under consideration, among other relevant factors (Karimi et al., 2020). Likewise, the long-term effects of fertilization regimes on soil microbial communities can vary according to the cropping system (e.g., legume vs. non-legume cropping systems) (Kong et al., 2023). In a meta-analysis of 37 studies reporting soil microbial diversity metrics in minerally-fertilized, organically-fertilized, and unfertilized soils, Bebber and Richards (2022) found that bacterial and archaeal taxonomic diversity was not significantly different between minerally-fertilized and unfertilized soil, but on average 2.9% greater in organically-fertilized vs. unfertilized soil, and 2.4% greater in organically-fertilized vs. minerally-fertilized soil. Nonetheless, they observed a very high residual heterogeneity in all meta-analyses of soil microbial diversity. Importantly, soil multifunctionality, i.e., its capacity to simultaneously sustain many functions (e.g., biomass production; source of raw materials; carbon pool; biodiversity pool; storing, filtering and transforming nutrients, substances and water) has been reported to increase with soil microbial diversity (Delgado-Baquerizo et al., 2016).

It is essential to identify and mechanistically understand the short-, medium- and long-term potential effects of agricultural practices on soil microbial diversity and, in general, on soil health and agroecosystem health. To this end, we must study the temporal dynamics in the evolution of agroecosystems, with special emphasis on the soil ecosystem as the basis of their functional integrity and sustainability. Given that long-term effects are often not only different from short-term effects but even contradictory, as well as the possibility of temporal concatenations of effects, it is crucial to carry out long-term field trials in which the changing nature, certainly not necessarily in a linear fashion, of the impacts derived from the application of agricultural practices and management systems can be observed.

Recent advances in metagenomic sequencing and computational tools (Karaoz et al., 2024), such as genome binning and metagenome-assembled genomes (MAGs) (Anantharama et al., 2016; Sharon and Banfield, 2013; Woodcroft et al., 2018), have considerably enhanced our capacity to explore soil microbial populations and communities, thereby fostering the understanding of their key roles for soil ecosystem functioning and functions. On the other hand, functional traits, defined as “morpho-physiophenological traits which impact fitness indirectly via their effects on growth, reproduction and survival, the three components of individual performance” (Violle et al., 2007), offer a promising avenue for linking taxonomic information to ecosystem functioning and functions. Thus, the deduction of ecologically-relevant microbial functional traits from the analysis of microbial genomes enables a trait-based assessment of microbial communities (Karaoz and Brodie, 2022). This opens the possibility to use functional trait ecology knowledge, initially developed especially for plants, to study microbial data and, hence, advance our understanding of microbial community assembly, species coexistence, biodiversity loss, etc., while enabling comparability across spatial and organizational scales (Dawson et al., 2021).

This study aimed to assess the permanence of the beneficial effects of the long-term (20 years) application of composted organic wastes, versus mineral fertilization or no fertilization, on vineyard soil functioning and functions under Mediterranean conditions. Many studies have been carried out to assess the long-term effects of fertilization regimes on soil health and microbial diversity, but, to our knowledge, there is only scarce information concerning the temporal permanence of those effects once fertilization has finished. To this purpose, four years after ceasing fertilization, a comprehensive analysis of soil physicochemical and biological properties, including prokaryotic diversity and functional traits by genome-centric metagenomics, was conducted to ascertain whether the beneficial effects of organic amendments were maintained 4 years after the end of fertilization. The genome-centric, trait-based analysis presented here provides a functional ecological assessment of the impact of the fertilization regimes on soil microorganisms (and, hence, soil health) by mapping microbial genomes to a trait space that covers energetic, resource acquisition, and stress tolerance traits which underlie ecologically-relevant microbial strategies associated with their environment. It was hypothesized that the beneficial effects of long-term organic management, in terms of soil health and multifunctionality, would persist after management had ended, due to the lag in re-equilibration of soil properties.

## Materials and methods

### Experimental design

The long-term field trial is located in the municipality of Bargota (province of Navarre, Spain), within the La Rioja Protected Designation of Origin. The climate is Mediterranean, with hot summers, an annual rainfall of 450-490 mm, and a mean annual temperature of 13.8 °C. The vineyard was planted in 1996 with Tempranillo variety, on a 5% slope in the catchment area of the Ebro River where the soil texture is loamy-clay and the soil is classified as a Calcixerept (Calleja-Cervantes et al., 2015).

The experiment followed a randomized block design with three replications. Each subplot had an area of 108 m^2^ and consisted of two rows of 18 m with 15 vines in each row. From 1998 to 2018, the following fertilization treatments were applied annually, except for the inorganic fertilization treatment which was applied biennially: (i) Abonlir, a pelletized compost made from plant and animal waste and sewage sludge produced by Slir S.L. at a rate of 3,700 kg FW ha^-1^ (Pellet treatment); (ii) a compost made from the organic fraction of municipal solid waste from Cárcar, Navarre at a rate of 4,075 kg FW ha^-1^ (Urban treatment); (iii) a compost made from sheep manure at a rate of 4,630 kg FW ha^-1^ (Manure treatment); (iv) an inorganic fertilization treatment [14 – 10 – 40 – 2.5 – 1.5 fertilizer units per hectare of nitrogen (N), phosphorus (P), potassium (K^+^), calcium (Ca^2+^), and magnesium (Mg^2+^), respectively] (Mineral treatment); and (v) a control treatment with no fertilization (Unfertilized treatment). The physical and chemical properties of these fertilizers are shown in Calleja-Cervantes et al. (2015). The rest of the agricultural management practices did not differ among treatments, e.g., chemical weed control each February, soil exposed during the vineyard cycle, ploughing the soil to a depth of 0-30 cm, and application of phytosanitary products). When necessary, a sprinkler irrigation system provided additional water. For more details on the experimental design, see Calleja-Cervantes et al. (2015).

### Soil sampling and analyses

Soil samples were taken in May 2022, four years after the last application of fertilizers. From each row within each subplot, composite (15 cores) soil samples were collected randomly at 0-10 cm depth with a core sampler. Samples were immediately taken to the laboratory, sieved to < 2 mm, and stored at 4 °C. Samples were then analysed for physicochemical and biological properties within one month. Soil particle size distribution, pH, cation exchange capacity (CEC), electrical conductivity (EC), dry matter content, nitrate and ammonium contents, Olsen phosphorus, active limestone, carbonates, and extractable Ca^2+^, Mg^2+^, Na^+^, and K^+^ were determined using standard methods (MAPA, 1994). Ceramic plates (Eijkelkamp, Giesbeek, The Netherlands) were used to measure both water-holding capacity at 33 kPa and wilting point. Total carbon (C) and N were measured using a Soli TOC Cube (Elementar) according to ISO 10694 (1995) and ISO 13878 (1998), respectively. Soil OM content was calculated by subtracting the carbonate content from the total C value and then multiplying the resulting organic C content by 1.72 (van Bemmelen, 1981). Mineral-associated OM (MAOM) and particulate OM (POM) were determined according to Cotrufo et al. (2019), together with soluble OM, using the 680 °C combustion catalytic oxidation method in a Shimadzu TOC-L analyzer (Kyoto, Japan). Soils were digested with a mixture of HNO_3_/HClO_4_ (Zhao et al., 1994) before the determination of the following elements by inductively coupled plasma atomic emission spectrometry (ICP-AES): aluminium (Al), arsenic (As), Ca^2+^, cadmium (Cd), cobalt (Co), copper (Cu), chromium (Cr), iron (Fe), lead (Pb), Mg^2+^, manganese (Mn), molybdenum (Mo), nickel (Ni), P, K^+^, sodium (Na^+^), sulphur (S), and zinc (Zn).

For soil biological properties, β-glucosidase, chitinase, L-alanine aminopeptidase, L-leucine aminopeptidase, phosphomonoesterase, and arylsulfatase enzyme activities were determined following ISO22939 (2010) with added fluorogenic substrates (4-methylumbelliferyl and 7-amino-4-methylcoumarin) in 96-well microplates, as described in Anza et al. (2019). Soil respiration was measured following ISO16072 (2002). Potentially mineralizable nitrogen (PMN) was measured according to Powers (1980). Microbial biomass carbon (MBC) was determined following Vance et al. (1987). Community-level physiological profiles (CLPPs) of cultivable heterotrophic bacteria were determined using Biolog Ecoplates^TM^ following Epelde et al. (2008).

Soil DNA was extracted from 0.25 g DW soil at least in triplicate (some samples required more extractions in order to achieve the minimum amount of DNA required for sequencing) using the Power Soil™ DNA Isolation Kit (MoBio Laboratories Inc., Carlsbad, CA). DNA quality and concentration were determined using a TapeStation^TM^ (Agilent Technologies Inc., Santa Clara, CA). Shotgun metagenomic sequencing, including library preparation, was performed at the Centro Nacional de Análisis Genómico (CNAG-CRG, Spain) using an Illumina NovaSeq6000 platform with a paired-end read length of 2 x 150 and yielding on average ∼18 +/-3.54 Gb of sequence data per sample (range: 11.21 - 25.83 Gb). Genomic data, including metagenome-assembled genomes and raw sequencing reads, are available under NCBI BioProject accession no. PRJNA1225280.

### Metagenome assembly, genome binning, and bin dereplication

Assembly of raw reads into scaffolds and binning scaffolds in genomes was done in KBase, available as a KBase narrative at https://kbase.us/n/126617/119/. Raw reads from each sample were imported into a KBase narrative (Arkin et al., 2018) and assembled into scaffolds using “JGI Metagenome Assembly Pipeline v0.6.0” (Kbase, 2025). Shortly, this workflow deploys the following steps: (1) quality-trimming and artifact removal with BBTools RQCFilter (Bushnell et al., 2017); (2) read correction with a Bloom filter (Li, 2015); (3) read pairing after filtering and correction with Seqtk; (4) assembly with metaSPAdes (default k-mers values of 33, 55, 77, 99, and 127) (Nurk et al., 2017); (5) BBTools stats.sh script for computation of assembly statistics; and (6) BBMap for mapping the reads to the scaffolds to quantify read representation in the scaffolds.

For each sample, the scaffolds were clustered into bins using MetaBAT2 (Kang et al., 2019) using coverage information from that sample and other samples from the same rowblock (6 binning groups with 5 samples each). The resulting bins from each sample were pooled to generate 1,918 bins where 861 bins had completeness > 50% and contamination < 25%. These bins were dereplicated using dRep at 99% ANI (Olm et al., 2017) (“completeness: 50, contamination: 25, secondary ANI threshold: 0.99, primary ANI threshold: 0.9) to yield 200 dereplicated bins (metagenome assembled genomes - MAGs). GTDB taxonomy (Parks et al., 2022) was assigned to MAGs using GTDB-Tk (Chaumeil et al., 2022).

### Quantification of MAG coverages across samples

MAG abundance profiles across samples were computed with coverM (Aroney et al., 2025) in “genome” mode (coverM genome) using dereplicated bins and raw reads from each sample as inputs. For each sample, coverM maps the reads to all the scaffolds from all these dereplicated bins, computes coverage statistics at scaffold level, and summarizes coverage metrics at genome level. The alignment thresholding options were set as “min-read-aligned-percent = 0.75, min-read-percent-identity = 0.95, min-covered-fraction-0”.

Mean coverages of genomes from samples were normalized for sequencing depth to generate a genome coverage table (200 genomes x 30 samples) (Table S1).

### Microbial community composition and diversity analysis

Prokaryotic community composition was assessed using single-copy ribosomal protein gene rpS3 chosen for its strong phylogenetic signal (Diamond et al., 2019; Hug et al., 2016). Open reading frames (ORFs) were predicted from contigs longer than 1 Kb using Prodigal V2.6.3 (Hyatt et al., 2010) in metagenome mode with the parameters “-m -p meta”. rpS3 marker sequences were identified across all metagenome assemblies using a hidden markov model (HMM) based on the alignment of rpS3 (rpsC) sequences from the tree of life data set from Hug et al. (2016). Sequences were clustered at 99% identity using USEARCH (Edgar, 2010) with the following options: “-cluster_fast -sort length -id 0.99-maxrejects 0-maxaccepts 0” to define species groups (SG). For each SG, the longest contig for the corresponding cluster was identified. Reads from each sample were mapped to these contigs using Bowtie2 in sensitive mode (--sensitive). The resulting bam files were used to generate a read count table using coverM contig command (version 0.5.0) [https://github.com/wwood/CoverM] with the following options: “--proper-pairs-only -- min-read-percent-identity 0.99 --min-read-aligned-percent 0.9 --methods count”. Normalized per base pair coverage for each contig was calculated with the following formula: reads mapped from sample / (contig length x reads sequenced in sample).

### Microbial fitness trait inference from MAGs

Microbial fitness traits were assigned to the dereplicated MAGs using microTrait, a HMM (hidden markov model) based workflow that maps genome sequences to a hierarchy of fitness traits based on the genomic basis of those traits from literature. The output from microTrait is a genomes x traits matrix with trait presence/absence (for binary traits) or count (for count traits) (Karaoz et al., 2022) (Table S2). For each trait and sample pair, sample level aggregated functional trait value was calculated by weighting trait presence/absence or counts by the normalized coverages of the genomes in the respective sample (Table S3).

### Statistical treatment

Soil multifunctionality was quantified as the average of multiple functions, previously standardized using a Z-transformation (Wagg et al., 2014). The soil properties used to estimate each of the investigated functions (i.e., water storage, fertility, carbon storage, and OM decomposition) were: (1) dry matter content, water-holding capacity, and wilting point for water storage; (2) total N, nitrate, ammonium, Olsen P, extractable Ca^2+^, extractable Mg^2+^, extractable K^+^, and sulfur content for fertility; (3) carbon content, MAOM, POM, and MBC for carbon storage; and (4) enzyme activities, respiration, PMN, and AWCD-Biolog^TM^ for OM decomposition. To assess differences among fertilization regimes on all measured soil parameters, as well as on diversity indices (Shannon, Simpson, Pielou), SGs, MAGs, and functional traits, linear mixed-effects models were used as implemented in the GAMLj module (Galluci, 2019) of the JAMOVI project (2022), with fertilization treatment as fixed effect and the subplot and row as random effects. Redundancy analyses (RDA) were performed using Canoco 5 (ter Braak and Šmilauer, 2012). Pheatmap and Rcmdr were used to draw clustered heatmaps and to calculate Spearman correlations between traits and soil parameters in R.

## Results

### Effects of fertilization regimes on soil properties

Many soil properties, both physicochemical and biological, showed statistically significant differences among fertilization treatments (Table 1 and Table S4), with significantly higher values being generally, though not always, observed in organically amended soils, compared to minerally-fertilized or unfertilized soils. However, despite such statistical significance, the quantitative magnitude of most of those differences was small. For example, soil pH was significantly higher in Pellet treatment, compared to all the other treatments, but the values ranged from 8.5 to 8.7. Cation exchange capacity was significantly higher in Manure treatment, compared to both Mineral and Unfertilized treatments, but the values varied from 13 to 14 mEq 100 g^-1^. Electrical conductivity was significantly highest in Pellet treatment. Dry matter content was significantly lower in Manure treatment, compared to both Mineral and Unfertilized treatments, with values fluctuating from 89 to 91%. Water-holding capacity was significantly lowest in Unfertilized treatment. Wilting point was significantly lower in Unfertilized treatment, compared to Urban and Manure, but values ranged from 13 to 15%. Organic matter, MAOM, and POM contents were significantly higher in the three organic treatments, compared to Mineral and Unfertilized treatments (similarly, soluble C content was significantly higher in Pellet treatment, compared to Mineral and Unfertilized treatments). Total N was significantly higher in Urban treatment, compared to Mineral and Unfertilized treatments. Carbonates content was significantly lower in Pellet treatment, compared to Mineral and Unfertilized treatments, with values ranging from 33 to 36%. Nitrate concentration was highest in Pellet treatment and lowest in Unfertilized treatment. Olsen P was significantly lower in Mineral and Unfertilized treatments than in the three organic treatments. Extractable Ca^2+^ and Mg^2+^ contents were significantly lowest and highest in Pellet and Manure treatment, respectively, although values varied from 26 to 28 mEq 100 g^-1^ for Ca^2+^ and from 1.5 to 1.8 mEq 100 g^-1^ for Mg^2+^. Significantly lower extractable Na^+^ values were observed in Pellet treatment, compared to Mineral and Unfertilized treatments, but values ranged from 0.11 to 0.15 mE 100 g^-1^. Potassium content was significantly higher in Pellet treatment than in Urban and Manure treatments, and significantly higher in these two organic treatments than in Mineral and Unfertilized treatments. Regarding heavy metals, significantly highest values of chromium, lead, and zinc were found in Urban treatment although, in terms of their quantitative magnitude, differences among treatments were relatively small. Sulfur content was significantly higher in the three organic treatments versus both Mineral and Unfertilized treatments. Finally, regarding enzyme activities, β-glucosidase and L-leucine aminopeptidase activities (as well as soil respiration) were significantly higher in Urban treatment, compared to Mineral and Unfertilized treatments. Conversely, arylsulfatase activity was significantly lower in the three organic treatments, compared to both Mineral and Unfertilized treatments.

**Table 1.** Effect of treatments on soil physicochemical and microbial properties (only those properties showing statistically significant differences among treatments are shown here; see also Supplementary Table 4). Values followed with different letters are significantly (p < 0.05) different. Mean values (n = 6) ± standard errors.

|  | Pellet |  | Urban |  | Manure |  | Mineral |  | Unfertilized |  |
| --- | --- | --- | --- | --- | --- | --- | --- | --- | --- | --- |
| Sand (%) | 23 ± 1 | a | 22 ± 2 | abc | 22 ± 1 | ab | 20 ± 1 | c | 21 ± 1 | bc |
| Cation exchange capacity (mEq 100 g<sup>-1</sup>) | 13 ± 0 | ab | 14 ± 1 | ab | 14 ± 1 | a | 13 ± 0 | b | 13 ± 0 | b |
| Electrical conductivity (ms cm<sup>-1</sup>) | 0.25 ± 0 | a | 0.21 ± 0 | bc | 0.20 ± 0 | bc | 0.21 ± 0 | b | 0.18 ± 0 | c |
| Dry matter content (%) | 90 ± 0 | ab | 90 ± 0 | ab | 89 ± 1 | a | 91 ± 0 | b | 91 ± 0 | b |
| Water-holding capacity at 33 kPa (%) | 27 ± 0 | a | 28 ± 1 | a | 28 ± 0 | a | 27 ± 1 | a | 25 ± 0 | b |
| Wilting point (%) | 14 ± 0 | ab | 14 ± 1 | a | 15 ± 1 | a | 14 ± 0 | a | 13 ± 0 | b |
| Organic matter (%) | 1.8 ± 0.1 | a | 2.0 ± 0.1 | a | 2.1 ± 0.1 | a | 1.3 ± 0.1 | b | 1.5 ± 0.1 | b |
| Total carbon (%) | 5.7 ± 0.1 | a | 6.0 ± 0 | b | 6.0 ± 0.1 | b | 5.7 ± 0.1 | a | 5.9 ± 0.1 | ab |
| Total nitrogen (%) | 0.14 ± 0 | ab | 0.17 ± 0 | c | 0.16 ± 0 | ac | 0.13 ± 0 | b | 0.14 ± 0 | ab |
| Mineral-associated organic carbon (mg g<sup>-1</sup>) | 8.7 ± 0.5 | a | 8.9 ± 0.6 | a | 8.5 ± 0.5 | a | 6.1 ± 0.2 | b | 6.4 ± 0.2 | b |
| Particulate organic carbon (mg g<sup>-1</sup>) | 2.2 ± 0.2 | a | 2.5 ± 0.2 | ab | 2.9 ± 0.2 | b | 1.5 ± 0.1 | c | 1.6 ± 0.1 | c |
| Extractable organic carbon (mg g<sup>-1</sup>) | 0.42 ± 0.1 | a | 0.29 ± 0 | ab | 0.28 ± 0 | b | 0.20 ± 0 | b | 0.21 ± 0 | b |
| Carbonates (%) | 33 ± 1 | a | 35 ± 1 | ab | 35 ± 1 | b | 36 ± 1 | b | 36 ± 0 | b |
| Nitrate (mg N-NO<sub>3</sub><sup>-</sup> kg<sup>-1</sup>) | 39 ± 2 | a | 32 ± 2 | b | 29 ± 1 | b | 30 ± 2 | b | 24 ± 1 | c |
| Olsen phosphorus (mg kg<sup>-1</sup>) | 79 ± 5 | ab | 85 ± 3 | a | 75 ± 3 | b | 46 ± 2 | c | 51 ± 4 | c |
| Extractable calcium (mEq 100 g<sup>-1</sup>) | 26 ± 0 | a | 28 ± 1 | b | 28 ± 0 | b | 28 ± 0 | b | 27 ± 0 | b |
| Extractable magnesium (mEq 100 g<sup>-1</sup>) | 1.5 ± 0 | a | 1.6 ± 0.1 | a | 1.8 ± 0.1 | b | 1.6 ± 0.1 | a | 1.6 ± 0.1 | a |
| Extractable sodium (mEq 100 g<sup>-1</sup>) | 0.11 ± 0 | a | 0.12 ± 0 | ab | 0.13 ± 0 | bc | 0.15 ± 0 | c | 0.13 ± 0 | b |
| Extractable potassium (mg kg<sup>-1</sup>) | 105<br>4 ± 49 | a | 785 ± 22 | b | 817 ± 26 | b | 656 ± 28 | c | 658 ± 55 | c |
| Cadmium (mg kg<sup>-1</sup>) | 0.27 ± 0 | a | 0.27 ± 0 | a | 0.25 ± 0 | ab | 0.25 ± 0 | ab | 0.23 ± 0 | b |
| Chromium (mg kg<sup>-1</sup>) | 24 ± 0 | a | 27 ± 2 | b | 24 ± 1 | a | 25 ± 1 | a | 23 ± 1 | a |
| Lead (mg kg<sup>-1</sup>) | 13 ± 0 | a | 19 ± 1 | b | 13 ± 1 | a | 13 ± 0 | a | 12 ± 1 | a |
| Magnesium (g kg<sup>-1</sup>) | 6.1 ± 0.1 | a | 6.3 ± 0.1 | b | 6.3 ± 0.1 | ab | 6.2 ± 0.1 | ab | 5.9 ± 0 | c |
| Phosphorus (g kg<sup>-1</sup>) | 1.62 ± 0.2 | a | 1.01 ± 0.1 | b | 0.89 ± 0.1 | bc | 0.65 ± 0 | c | 0.69 ± 0.1 | bc |
| Sodium (g kg<sup>-1</sup>) | 0.45 ± 0 | a | 0.39 ± 0 | b | 0.37 ± 0 | bc | 0.38 ± 0 | bc | 0.35 ± 0 | c |
| Sulfur (mg kg<sup>-1</sup>) | 0.64 ± 0 | a | 0.69 ± 0 | a | 0.67 ± 0 | a | 0.56 ± 0 | b | 0.57 ± 0 | b |
| Zinc (mg kg<sup>-1</sup>) | 57 ± 2 | a | 65 ± 3 | b | 50 ± 2 | c | 47 ± 1 | c | 51 ± 2 | c |
| β-glucosidase ($\mu\text{mol kg}^{-1} \text{ h}^{-1}$) | 698 ± 39 | a | 849 ± 44 | b | 462 ± 54 | c | 587 ± 55 | ac | 637 ± 46 | a |
| L-leucine aminopeptidase ($\mu\text{mol kg}^{-1} \text{ h}^{-1}$) | 388 ± 23 | a | 490 ± 22 | b | 339 ± 6 | ac | 312 ± 45 | c | 343 ± 26 | ac |
| Arylsulfatase ($\mu\text{mol kg}^{-1} \text{ h}^{-1}$) | 87 ± 11 | a | 70 ± 6 | ab | 29 ± 10 | b | 161 ± 21 | c | 145 ± 20 | c |
| Respiration ($\text{mg C-CO}_2 \text{ kg}^{-1} \text{ h}^{-1}$) | 0.75 ± 0.1 | ab | 0.79 ± 0 | a | 0.80 ± 0 | a | 0.67 ± 0 | bc | 0.58 ± 0 | c |

These results show that, while statistically significant differences were observed among fertilization treatments, generally, though not always, with higher values in the organic treatments, the magnitude of the differences was small. But, when individual soil properties were grouped into soil functions (i.e., water storage, fertility, carbon storage, OM decomposition), the average Z-scores for all four functions, as well as for the multifunctionality index, were mostly positive for Manure, Urban, and Pellet treatments, and negative for Mineral and Unfertilized treatments (differences were statistically significant in many cases) (Figure 1). An exception was the OM decomposition function in Manure treatment, which showed a negative value.

**Figure 1.**
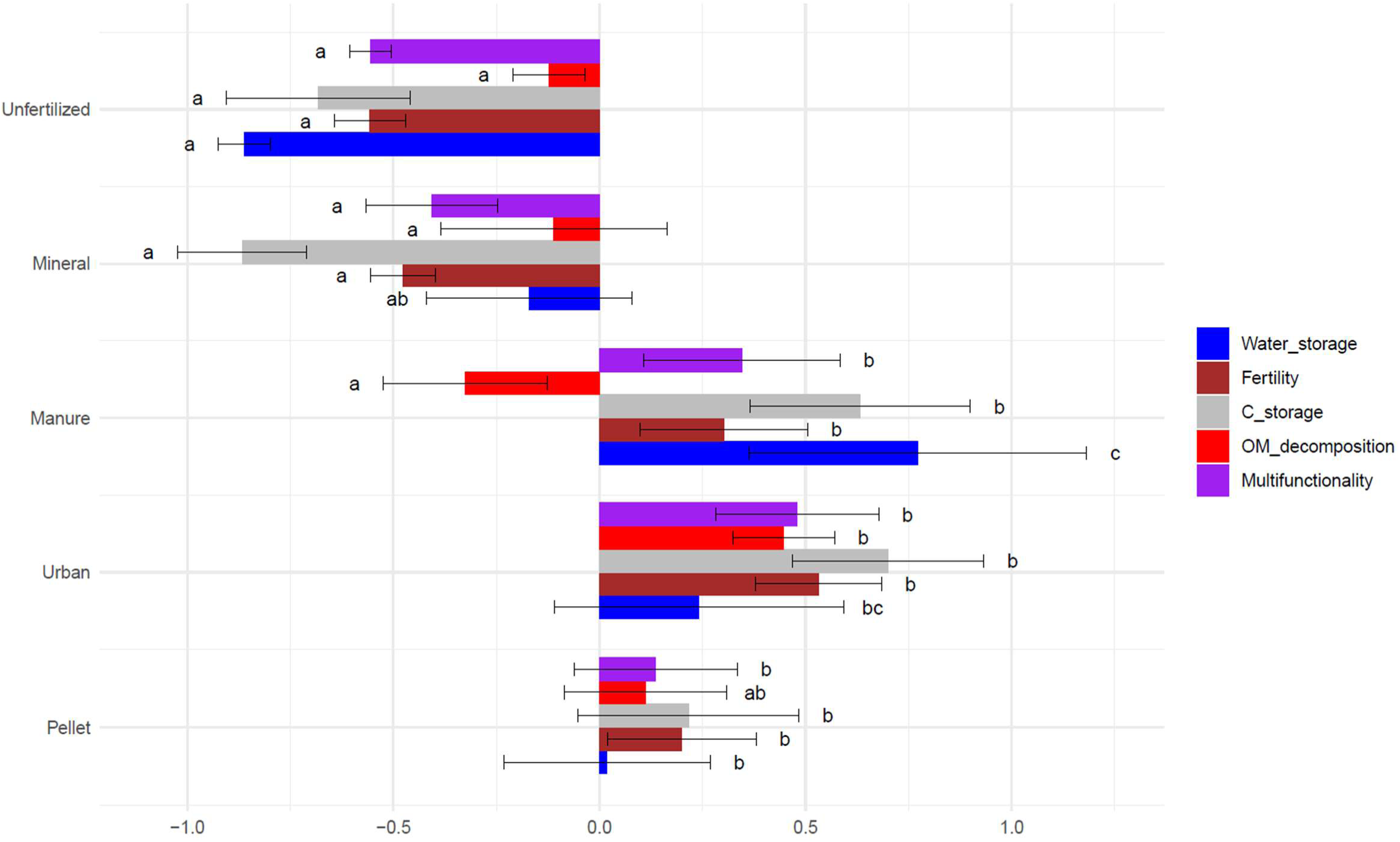
Z-scores of the soil properties related to each soil function and then averaged to obtain a multifunctionality index. Means (n = 6) ± standard errors. Bars followed with different letters indicate significant differences among treatments (P < 0.05).

### Effect of fertilization regimes on soil prokaryotic structural diversity

From the soil prokaryotic diversity analysis, 600 species groups (SGs) were obtained. For structural diversity indices calculated from SGs (Table 2), statistically significant differences were observed, but clear trends were lacking and the magnitude of differences among treatments was small. The Unfertilized treatment showed significantly higher values for a number of diversity indices, including the (i) Shannon index, compared to Mineral and Urban treatments; (ii) Simpson index, compared to the three organic treatments; and (iii) Pielou index, compared to the Pellet and Urban treatments.

**Table 2.** Effect of treatments on structural (top) and functional (bottom) diversity indices calculated from the species groups and traits predicted by microTrait, respectively. Means (n = 6) ± standard errors. Values with different letters are significantly different (P < 0.05).

| STRUCTURAL |  |  |  |
| --- | --- | --- | --- |
|  | Shannon | Simpson | Pielou |
| Pellet | 6.021 ± 0.008<sup>ab</sup> | 0.99640 ± 0.00011<sup>a</sup> | 0.9482 ± 0.0011<sup>a</sup> |
| Urban | 6.001 ± 0.018<sup>a</sup> | 0.99650 ± 0.00008<sup>a</sup> | 0.9501 ± 0.0010<sup>a</sup> |
| Manure | 6.019 ± 0.018<sup>ab</sup> | 0.99662 ± 0.00008<sup>ab</sup> | 0.9505 ± 0.0014<sup>ab</sup> |
| Mineral | 6.008 ± 0.008<sup>a</sup> | 0.99673 ± 0.00005<sup>bc</sup> | 0.9507 ± 0.0013<sup>ab</sup> |
| Unfertilized | 6.054 ± 0.009<sup>b</sup> | 0.99685 ± 0.00004<sup>c</sup> | 0.9539 ± 0.0011<sup>b</sup> |
| FUNCTIONAL |  |  |  |
| Pellet | 3.658 ± 0.001<sup>a</sup> | 0.9355 ± 0.0001<sup>a</sup> | 0.749 ± 0.000<sup>a</sup> |
| Urban | 3.655 ± 0.002<sup>a</sup> | 0.9350 ± 0.0001<sup>b</sup> | 0.748 ± 0.000<sup>a</sup> |
| Manure | 3.652 ± 0.002<sup>a</sup> | 0.9346 ± 0.0002<sup>c</sup> | 0.748 ± 0.000<sup>a</sup> |
| Mineral | 3.664 ± 0.003<sup>b</sup> | 0.9356 ± 0.0002<sup>a</sup> | 0.750 ± 0.001<sup>b</sup> |
| Unfertilized | 3.654 ± 0.002<sup>a</sup> | 0.9350 ± 0.0001<sup>bc</sup> | 0.748 ± 0.000<sup>a</sup> |

Fertilization treatments had a significant influence on the structural composition of soil prokaryotic communities, accounting for 18% of the variation in the data (pseudo-F = 1.4, p = 0.002). As shown in the RDA biplot (Figure 2), Unfertilized and Mineral treatments separated from the three organic treatments along the horizontal axis, while Pellet treatment separated from Urban and Manure treatments along the vertical axis. The biplot shown in Figure 2 only includes the 20 SGs with the best fit that influenced the observed differences among fertilization treatments, but other SGs also showed statistically significant differences among treatments (Table S5).

**Figure 2.**
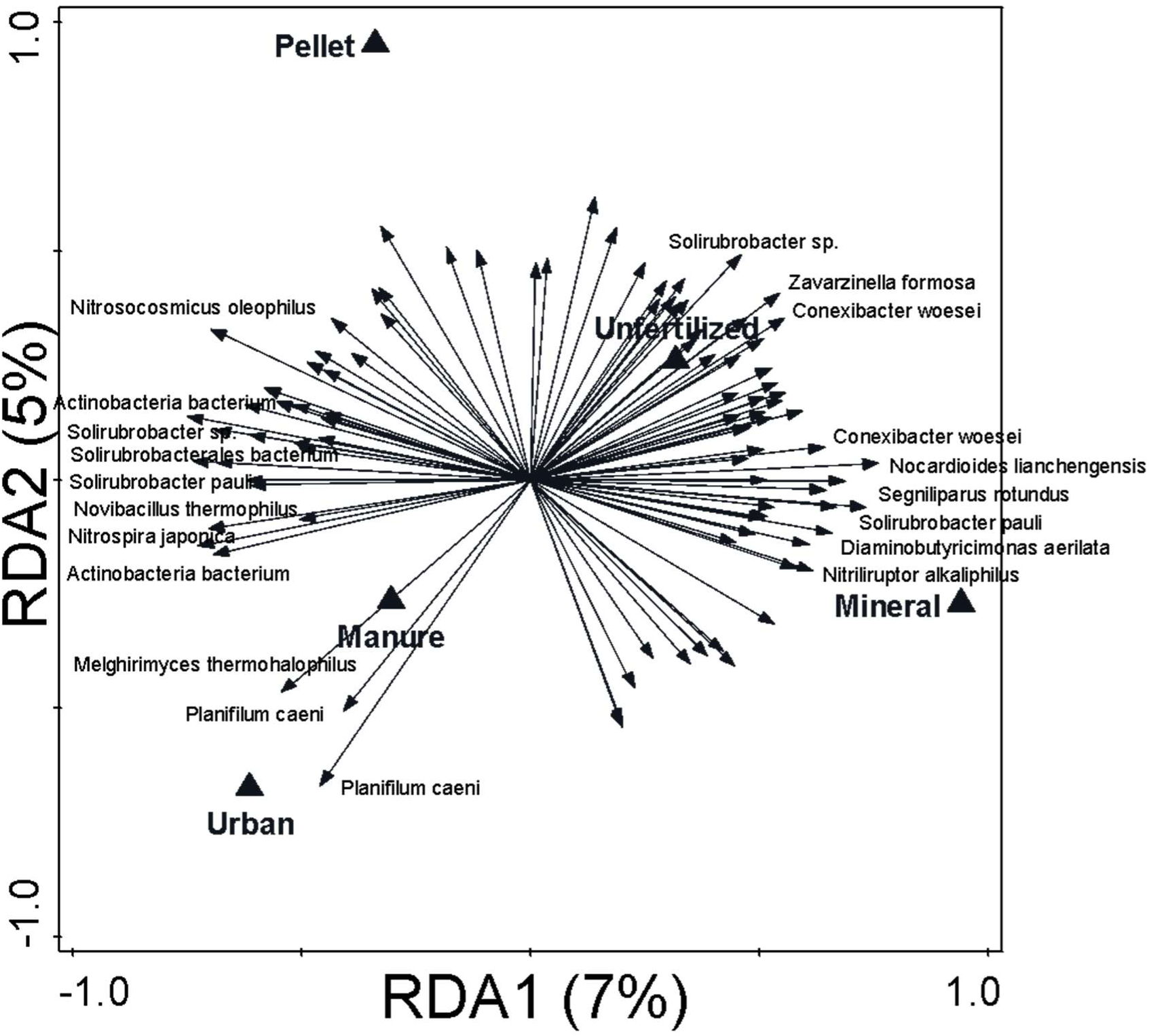
Biplot of the redundancy analysis (RDA) performed on species groups data as response variable and applied treatments as explanatory variables. Only the best-fitting 20 taxa have been labelled. Variance explained by each axis is shown between brackets.

Twenty-one SGs with statistically significant differences among treatments showed higher abundance values in the organic treatments, compared to Mineral and Unfertilized treatments (interestingly, two taxa, i.e. *Nitrosocosmicus oleophilus* and *Nitrospira japonica*, showed very high abundances in the organic treatments) (Figure S2A). Conversely, 49 SGs with statistically significant differences among treatments showed lower abundance values in the organic treatments (Figure S2B), with SG assigned to *Segniliparus rotundus* and *Diaminobutyricimonas aerilata* being particularly abundant in Unfertilized and Mineral treatments. There were only two SGs whose abundance in organic treatments was higher than in Mineral treatment but lower than in Unfertilized control (Figure S2C). By contrast, there were 39 SGs in which the abundances of the organic treatments differed from each other when compared to Mineral and Unfertilized treatments (Figure S2D). In summary, out of the 600 SGs obtained here (100%), no significant differences were observed for 489 SGs (81.5%). Among the remaining 111 SGs (those showing statistically significant differences), 21 and 49 SGs showed higher and lower abundance values, respectively, in the organic treatments, compared to Mineral and Unfertilized treatments.

Two hundred MAGs were obtained in this study. The RDA biplot in Figure S3 shows a very similar pattern to that obtained from SGs (Figure 2), in this case explaining 45% of the variation in the data (pseudo-F = 5.2, p = 0.002). Again, Unfertilized and Mineral treatments separated from the three organic treatments along the horizontal axis, while Pellet treatment separated from Urban and Manure along the vertical axis. Table S5 shows that 70% of the MAGs showed statistically significant differences among treatments.

### Effect of fertilization regimes on soil prokaryotic functional diversity

The microTrait workflow allowed the assignment of microbial fitness traits to the MAGs. Concerning functional diversity indices (Table 2), the Mineral treatments showed higher values of the Shannon and Pielou indices, compared to all the other treatments, and higher values of the Simpson index compared to all but the Pellet treatment.

In terms of resource use traits, organic amendments promoted the proliferation of chemotrophs as opposed to phototrophs (Figure 3). There was a significantly higher abundance of chemoorganoheterotrophs in Urban and Manure treatments, chemolithoheterotrophs in Pellet and Mineral treatments, and chemolithoautotrophs in Pellet treatment, compared to Unfertilized treatment. However, significantly higher and lower abundances of photoautotrophs were found in Mineral and Urban treatments, respectively. Mineral treatment led to statistically significant higher values of most resource acquisition traits (S, N, C1 and P compounds acquisition, substrate uptake and simple compound degradation), compared to Unfertilized treatment. Similarly, desiccation/osmotic/salt stress, pH stress, oxidative stress, temperature tolerance, and general stress tolerance showed higher values in Mineral treatment. Stress tolerance traits tended to show lower values in organic treatments, while oxygen limitation and prokaryotic envelope stress tended to show higher values.

**Figure 3.**
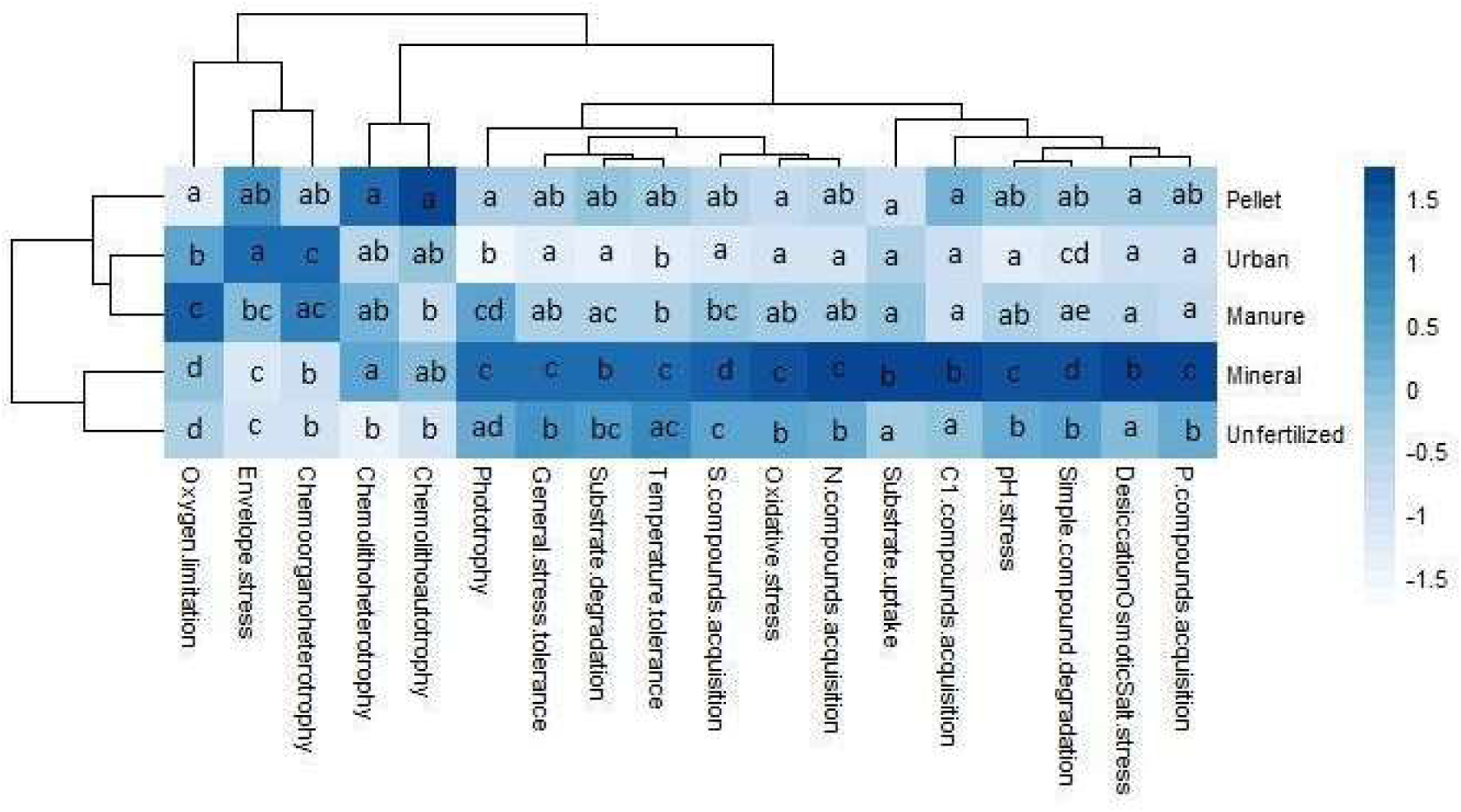
Heatplot of the average relative abundance of traits at second level of hierarchy. Values followed with different letters are significantly different among treatments (P < 0.05).

The distribution of soil samples in functional terms was very similar to that observed in structural terms. The RDA in Figure 4 explains 34% of the variation in the data (pseudo-F = 3.2, p = 0.004). Mineral and Unfertilized treatments appear separated from the three organic treatments along the horizontal axis, which explains most of the variability. Most traits that shower higher values in Mineral treatment were those related to resource acquisition, more specifically, traits related to substrate assimilation. The Unfertilized treatment showed higher values of traits related to phototrophy (i.e., isorenieratene, betaisorenieratene, chlorobactene). The different organic treatments appear separated among themselves along the vertical axis. Table S3 shows the normalized coverage of traits for each sample, where up to 86% of the traits showed statistically significant differences among treatments.

**Figure 4.**
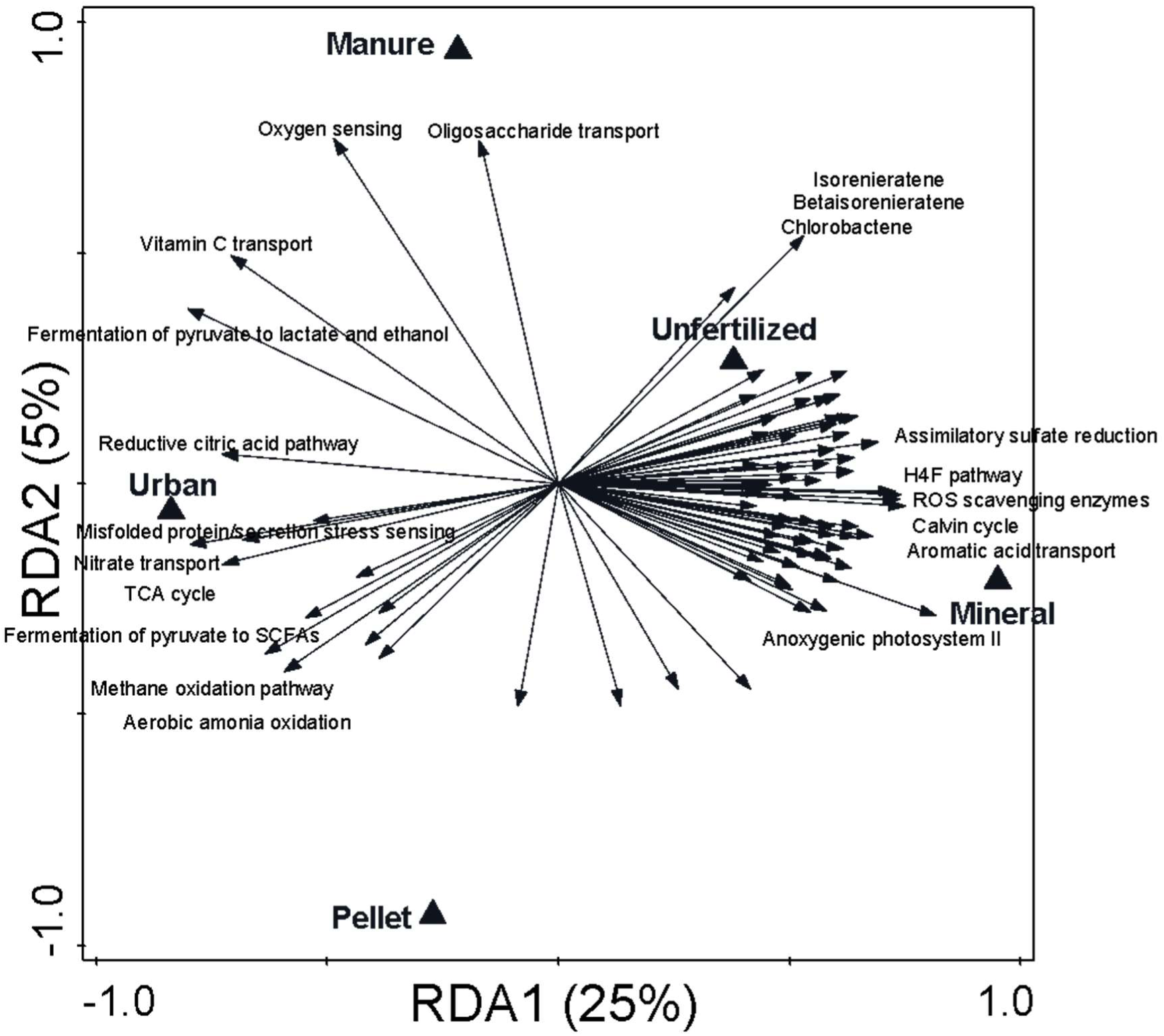
Biplot of the redundancy analysis (RDA) performed on trait data at third level of hierarchy as response variable and applied treatments as explanatory variables. Only the best-fitting 20 traits have been labelled. Variance explained by each axis is shown between brackets.

### Relationship between soil functions and prokaryotic functional traits

Prokaryotic envelope stress was highly positively correlated with all soil functions, as well as with the values of the multifunctionality index (Figure 5). Prokaryotic envelope stress was also positively correlated with Cd, Mn, Pb, Zn, Olsen P, MAOM, and OM contents (Figure S4). At the third hierarchical level (Table ST6), nine of the thirteen genes that showed significant positive correlations with soil functions were associated with key processes in the biogeochemical cycles of nitrogen and carbon. Specifically, nitrogen-related genes were involved in nitrate and nitrite reduction, nitrogen and urea transport, the urease system, and aerobic ammonia oxidation, while carbon-related genes were linked to cell membrane lipid transport, the reductive citric acid pathway, methane oxidation, and the fermentation of pyruvate to short-chain fatty acids (SCFAs).

**Figure 5.**
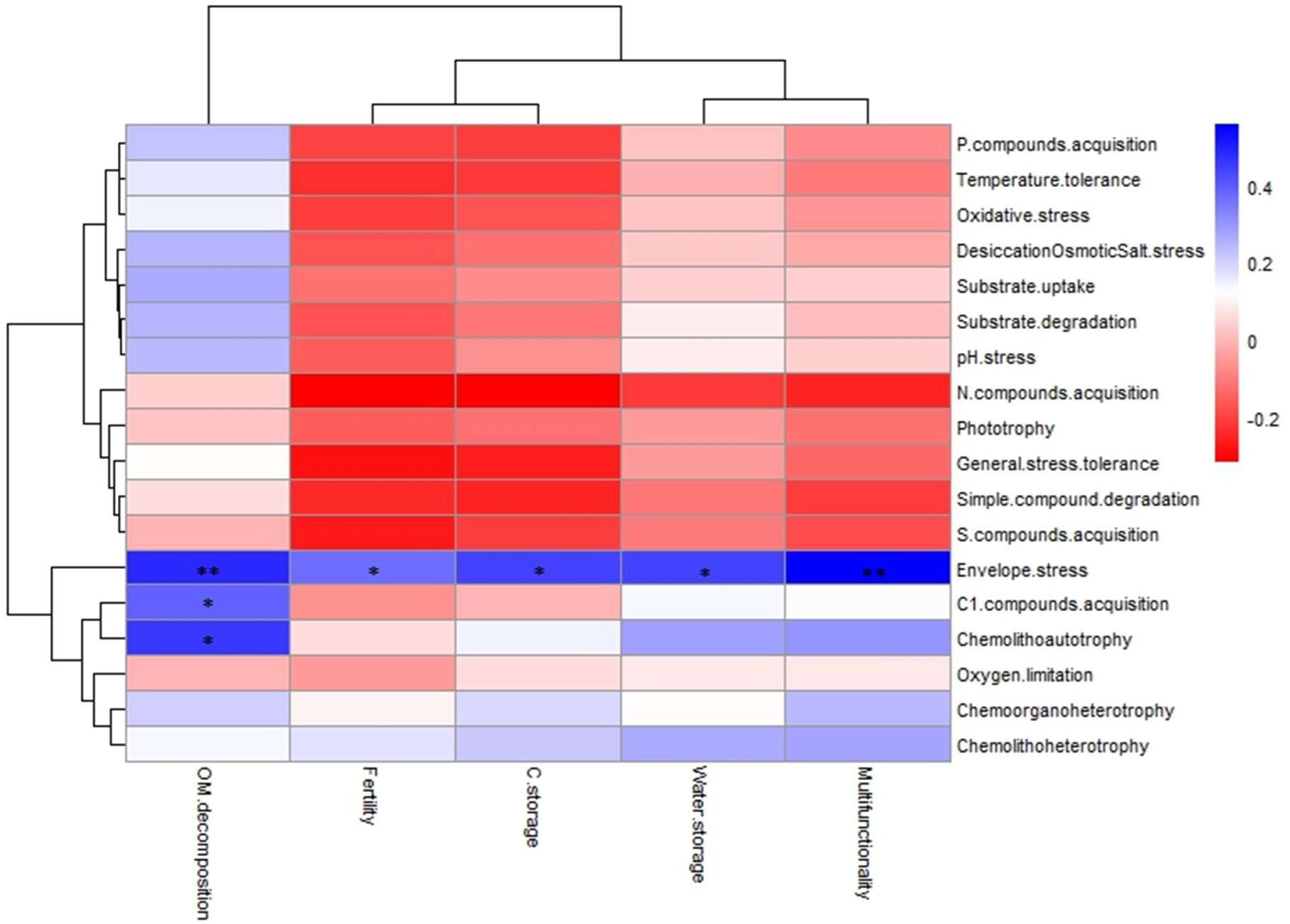
Heat map showing the Spearman correlations between traits at second level of hierarchy and soil functions (*: 0.01 < p ≤ 0.05; **: 0.001 < p ≤ 0.01; ***: p ≤ 0.0001; n = 30).

## Discussion

The main objective of this study was to assess the permanence of the beneficial effects of the long-term (20 years) application of composted organic wastes, versus mineral fertilization or no fertilization, on vineyard soil functioning and functions under Mediterranean conditions. As indicated above, in general, but not always, soils treated with composted organic wastes showed significantly higher values of many soil properties, specifically those linked to the content of OM and nutrients (e.g., OM, MAOM, POAM, sulfur, Olsen P, extractable K^+^, etc.) (Table 1). However, despite such statistical significance, the quantitative magnitude of many of those differences was small. In a previous study on the same experimental assay, Calleja-Cervantes et al. (2015) also found significantly higher values of key soil properties in the organically-amended soils: compared to Mineral and Unfertilized treatments, they observed significantly higher values of (i) OM content in Manure treatment; (ii) N content in Pellet and Manure treatments; (iii) Olsen P content in the three organic treatments; and (iv) K^+^ content in Manure treatment. Therefore, it was concluded that, four years after ceasing fertilization, the beneficial effect of the long-term (20 years) application of composted organic wastes on those key soil properties is still present.

By contrast, arylsulfatase activity (it catalyses the hydrolysis of organic sulfate esters to sulfate-S, thus reflecting S turnover and cycling in soils) was significantly lower in the three organic treatments. This could be due to the significantly higher S contents found in the organically-amended soils (in other words, due to feedback inhibition by available inorganic S). Mocali et al. (2020) suggested that S cycling can play a significant role in grapevine plant health and wine quality, since a reduced availability of sulfates for vine plants might limit the reduced glutathione synthesis (reduced glutathione plays a key role in aroma protection in musts and wines). Nonetheless, compared to those enzymes involved in soil C, N, and P cycling, arylsulfatase activity and its regulation (e.g., its controlling factors) has been much less investigated (Chen et al., 2019).

Finally, regarding heavy metals, significantly highest values of chromium, lead, and zinc were found in Urban treatment although, in terms of their quantitative magnitude, differences among treatments were relatively small. While the observed soil metal concentrations are not currently alarming, the presence of metals in the organic amendments must always be taken into consideration when recommending their use, as their prolonged application may lead to toxicologically-relevant soil and plant metal accumulation and, hence, potential hazards for ecosystems and humans (Déportes et al., 1998).

Most importantly, the grouping of soil properties into soil functions, as well as the calculation of an index of soil multifunctionality, also point to an enhanced soil health in the three organically-fertilized soils. The use of organic amendments has often been associated with enhanced soil properties (Urra et al., 2019), while fostering beneficial microorganisms (Bulluck et al., 2002). In particular, the C content of vineyard soils affects soil fertility, erosion, and biogeochemical cycles, with important repercussions for the global climate (Giffard et al., 2022). On the other hand, in the absence of organic amendments, intensive viticulture often decreases topsoil organic C (Jakšić et al., 2020). Importantly, the application of organic amendments provides a direct source of C for soil microorganisms, as well as an indirect one via increased crop growth and plant residue returns (Bünemann et al., 2006). In the long-standing debate over concepts such as soil health, quality, fertility, functions, capability, and ecosystem services (Bünemann et al., 2018), it is important to emphasize that ‘more’ does not always mean ‘better’ when evaluating the effects of agricultural practices such as organic farming on soil functions and multifunctionality. Potential trade-offs and synergies among functions must be considered, and outcomes are strongly influenced by the soil properties and processes selected for function calculation, as well as by overlapping indicators (e.g., soil OM) (Zhao et al., 2022). Zwetsloot et al. (2021) proposed that most agricultural fields can sustain at least three soil functions at optimal levels before trade-offs emerge. Because soil functions arise from complex interactions among physical, chemical, and biological processes, caution is needed when interpreting the effects of management on soil functionality. Moving beyond the traditional minimum dataset approach, Debeljak et al. (2019) applied multi-criteria decision models to assess soil functional capacity, decomposing complex functions into subfunctions to evaluate overall multifunctionality. Creamer et al. (2022) further developed a framework to quantify and communicate the role of soil biology in driving multifunctionality.

Regarding prokaryotic structural diversity reflected by metagenomic data, it was observed that the three organic amendments significantly affected microbial community composition. However, previous studies report inconsistent effects of organic versus mineral fertilization on soil microbes (Cui et al., 2023; Siles et al., 2024; Hendgen et al., 2018; Ali et al., 2022; Durrer et al., 2021; Mo et al., 2020; Shah and Wu, 2020). Such variability likely arises from factors like soil pH, texture, crop type, amendment dose, and environmental conditions (Cui et al., 2023), a fact which might, at least partly, explain the abovementioned inconsistencies and, hence, unpredictable results. There is certainly a need for more in-depth studies to better understand and predict the magnitude and direction of the responses of soil microbial communities to the application of organic amendments. In this respect, long-term studies are of great importance since, given the dynamic nature of the soil ecosystem, it is almost inevitable that the effects of organic amendments on soil microbial communities will change over time. These changes will probably be influenced by the microbeś life history strategies: for instance, oligotrophic and copiotrophic microorganisms (Fierer et al., 2012) will exhibit different carbon-use efficiencies with concomitant consequences for OM decomposition and mineralization, and hence microbial structure.

Methodological differences likely also explain the observed discrepancies. High-throughput sequencing has greatly advanced the study of soil microbial communities (Chen et al., 2022; Singer et al., 2016), including their responses to organic amendments (Li et al., 2022a, 2022b; Zhang et al., 2021, 2022a, 2022b). 16S rRNA amplicon sequencing is commonly used for taxonomic and phylogenetic analyses, and this was the method applied in the same experimental field by Calleja-Cervantes et al. (2015). Meanwhile, we assessed soil prokaryotic community composition using the single-copy ribosomal protein gene rpS3, which provides a stronger phylogenetic signal and is more reliably reconstructed from metagenomic data (Diamond et al., 2019; Hug et al., 2016; Sharon et al., 2015; Miller et al., 2011; Hug et al., 2013). Compared to 16S-based approaches, shotgun metagenomics and rpS3 analysis offer improved detection of bacterial diversity, enhanced gene prediction, and more robust phylogenetic resolution (Ranjan et al., 2016; Probst et al., 2018). With this approach, in our study, it was observed that organic treatments fostered the proliferation of specific taxa, such as *Nitrosocosmicus oleophilus* (ammonia-oxidizing archaea) and *Nitrospira japonica* (nitrite-oxidizing bacteria), both linked to nitrification, a two-step process where ammonia is oxidized to nitrite by ammonia-oxidizing bacteria and/or archaea, and subsequently to nitrate by nitrite-oxidizing bacteria (van Kessel et al., 2015). In our study, MAGs from the Nitrososphaeraceae family, particularly Nitrososphaerales archaeon TH5893, also increased in organically-amended soils. The amoA gene was not detected in the TH5893 MAG (Sheridan et al., 2020); therefore, its metabolic lifestyle remains unresolved, though it may possess mixotrophic potential and utilize organic nitrogen from the organic amendments. Nitrososphaeraceae has been shown to respond to soil N levels: in agricultural plots intensively fertilized with N, Nelkner et al. (2019) found that the four most abundant MAGs in bulk soil were related to the Nitrososphaeraceae family. Orellana et al. (2018) found that six deep-branching Thaumarchaeota and three complete ammonia oxidizer (comammox) Nitrospira populations increased in abundance up to 5-fold with N fertilizer addition, indicating that they respond rapidly to N fertilization. In our study, total N was only significantly higher in Urban treatment, compared to Mineral and Unfertilized treatments (no significant different among treatments were observed for soil ammonium content or PMN).

Concerning structural diversity indices calculated from SGs (Table 2), fertilization produced only minor effects on structural diversity, with the fertilized treatments showing slightly higher Shannon, Simpson, and Pielou indices than the unfertilized. A priori, one would expect that organic amendments would result in a positive effect on the abundance of microbial taxa characterized by a copiotrophic strategy but, due to the complexity of the soil ecosystem and, in particular, the complex relationships between the dynamics of microbial community composition in soil and C turnover in agroecosystems, that is not always the case. In a long-term soil restoration experiment carried out under semi-arid conditions, Siles et al. (2024) found that organic amendments reduced soil prokaryotic alpha diversity. In contrasts, Hendgen et al. (2018) found that organic management applied for 10 years yielded higher soil bacterial richness, compared to integrated management, in a German vineyard. Nonetheless, in their study (Hendgen et al., 2018), soil bacterial communities may have been positively affected by the reduced use of pesticides characteristic of organic farming. It is not uncommon to find land managers that assume that the application of organic amendments always increases soil microbial diversity. This is certainly not the case (in fact, it is common to find the opposite effect), just as it is not correct to assume that the more soil biodiversity the healthier the soil. It is true that a higher biodiversity may imply higher functional capabilities, with potential concomitant effects on soil health and multifunctionality, but up to a certain point (Grządziel, 2017).

The functional roles of microbial species are paramount for sustainable agriculture, especially those roles affecting soil fertility and crop productivity. The functional trait concept, especially when complemented with taxonomic information, provides critical insights into species’ roles in ecosystems, facilitating a transition from species identity to ecosystem function (Eisenhauer et al., 2022). Besides, in terms of the often-mentioned functional redundancy in soil, it must be stated that, although the basic general processes (e.g., respiration, C turnover, OM decomposition) are usually not affected by the loss of microbial richness, rare functions performed by only one or two specific groups of microorganisms (i.e., a narrow niche function) can indeed be impaired by a reduction in biodiversity. An example of such rare functions is the activity of nitrifying taxa, that are phylogenetically constrained across the tree of life and which were promoted by the organic amendments in this study.

In relation to functional diversity indices (Table 2), the Mineral treatments showed higher values of the Shannon and Pielou indices, compared to all the other treatments, and higher values of the Simpson index compared to all but the Pellet treatment. In functional terms, the Mineral treatment also led to significantly higher values of resource acquisition traits, including substrate assimilation and degradation, as well as tolerance traits to various stressors such as desiccation, osmotic/salt stress, pH stress, and oxidative stress. This fact likely contributed to the elevated functional diversity observed in the Mineral treatment. A plausible hypothesis is that the Mineral treated soil might lack some important macro- or micro-nutrients present in organically-treated soils, resulting in resource limitation and stress, fostering microbial communities characterized by a combination of resource acquisition and stress tolerance strategies (Malik et al., 2020). However, such metabolic investments may reduce cellular growth efficiency and lead to lower biomass values. Nonetheless, in our study, no significant differences were observed between organically-fertilized and minerally-fertilized soils in terms of MBC (some statistically significant differences were observed for soil respiration values but not for PMN).

Organic treatments promoted the proliferation of chemotrophic communities, possibly due to the added compounds present in the amendments. Stress tolerance traits decreased with organic fertilization, likely due to the improved soil properties and functions. Organic treatments created conditions more amenable to organisms that can deal with oxygen limitation and induced prokaryotic envelope stress. The envelope stress response is triggered to maintain cellular homeostasis when environmental conditions may cause damage (Mitchell and Silhavy, 2019). The increased water storage capacity of organically-fertilized soils may have induced oxygen limitation, as this trait was positively correlated with the water storage function.

Finally, the study of the relationships between soil functions and microbial functional traits revealed that most traits were negatively correlated with soil functions, suggesting that healthier soils with higher soil OM contents result in fewer stresses on soil microbial communities, reducing their reliance on resource acquisition traits. Also, the significant positive correlations between soil functions and genes related to nitrogen transformations (e.g., urease activity, ammonia oxidation, nitrate and nitrite reduction) indicate that active nitrogen turnover may be a key driver of soil biochemical processes, possibly enhancing nutrient availability and promoting microbial productivity. Similarly, the association of soil functions with carbon-related genes such as methane oxidation, pyruvate fermentation to SCFAs, and the reductive citric acid pathway highlights the contribution of carbon fluxes to energy generation and organic matter turnover within the microbial community. Together, these findings point to a close coupling of nitrogen and carbon metabolic pathways in regulating soil multifunctionality, suggesting that shifts in the abundance of these genes could serve as sensitive indicators of soil health and resilience to environmental change.

In conclusion, our study demonstrates that the long-term application of organic amendments provides lasting benefits to vineyard soil health and functionality, even four years after application ceased. The beneficial effects of organic amendments on vineyard soil are key for sustainable wine production, with potential implications for terroir and grape quality.

## CRediT authorship contribution statement

Lur Epelde: Conceptualization, formal analysis, funding acquisition, writing – original draft. Ulas Karaoz: Conceptualization, formal analysis, writing – review and editing. Carlos Garbisu: Conceptualization, funding acquisition, writing – review and editing. José Félix Cibriáin: Conceptualization, funding acquisition, writing – review and editing. Eoin Brodie: Conceptualization, writing – review and editing.

## Declaration of competing interest

The authors declare that they have no known competing financial interests or personal relationships that could have appeared to influence the work reported in this paper.

## Funding sources

This research has been funded by (i) the Ikermugikortasuna mobility programme of the Basque Government, (ii) the grant ReCROP PCI2021-121935 funded by MICIU/AEI/10.13039/501100011033 and the European Union Next Generation EU/PRTR, (iii) the European Union’s Horizon Europe research and innovation action under grant agreement No. 101086179 (AI4SoilHealth), and (iv) the grant VinAE PCI2025-163141 funded by MICIU/AEI/10.13039/501100011033.

---

**Supplementary Figure 1.**
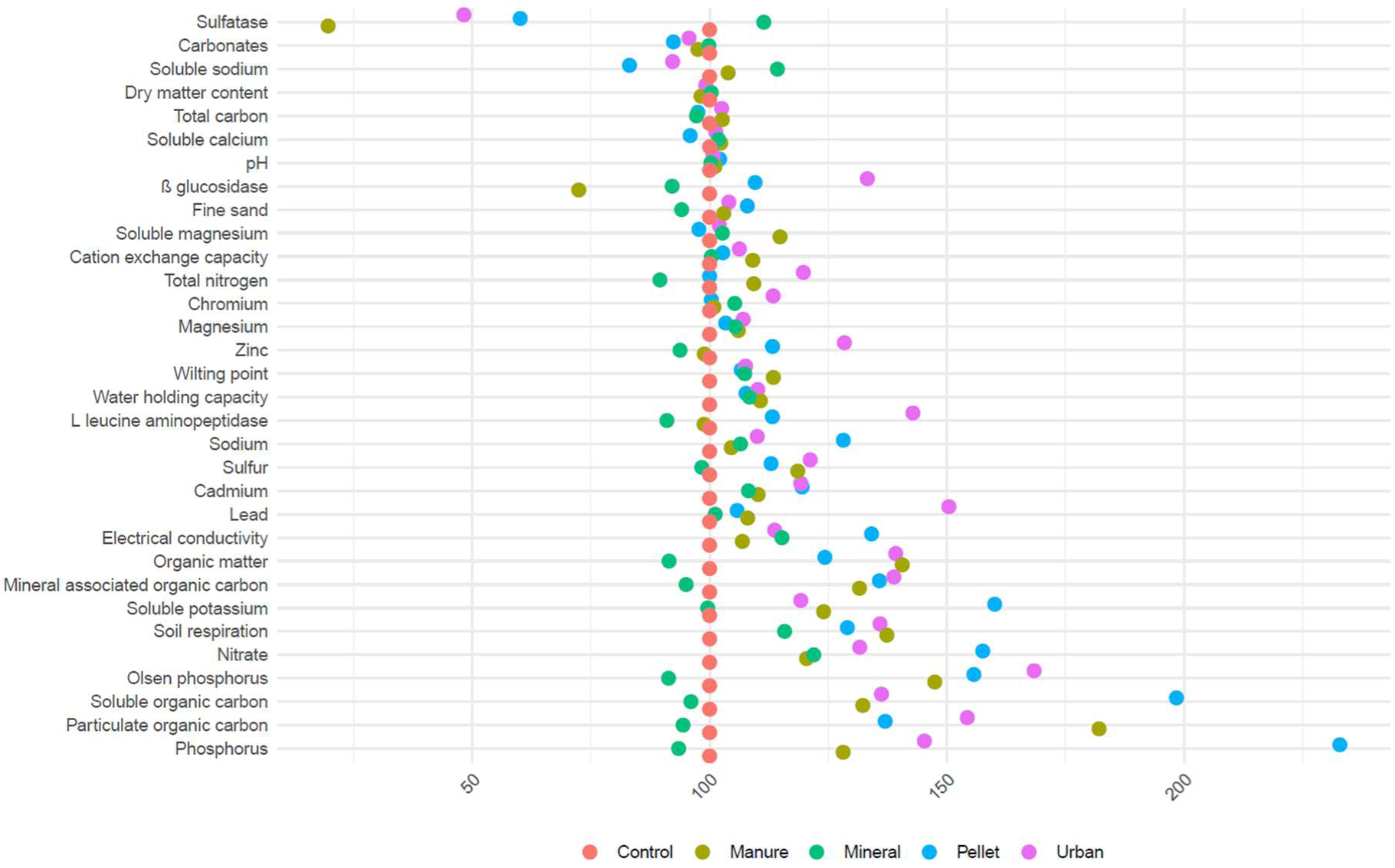
Dotplot showing the changes relative to Unfertilized treatment (control) in those soil properties with statistically significant differences. Soil properties are arranged from bottom to top by their increase relative to Unfertilized treatment.

**Supplementary Figure 2.**
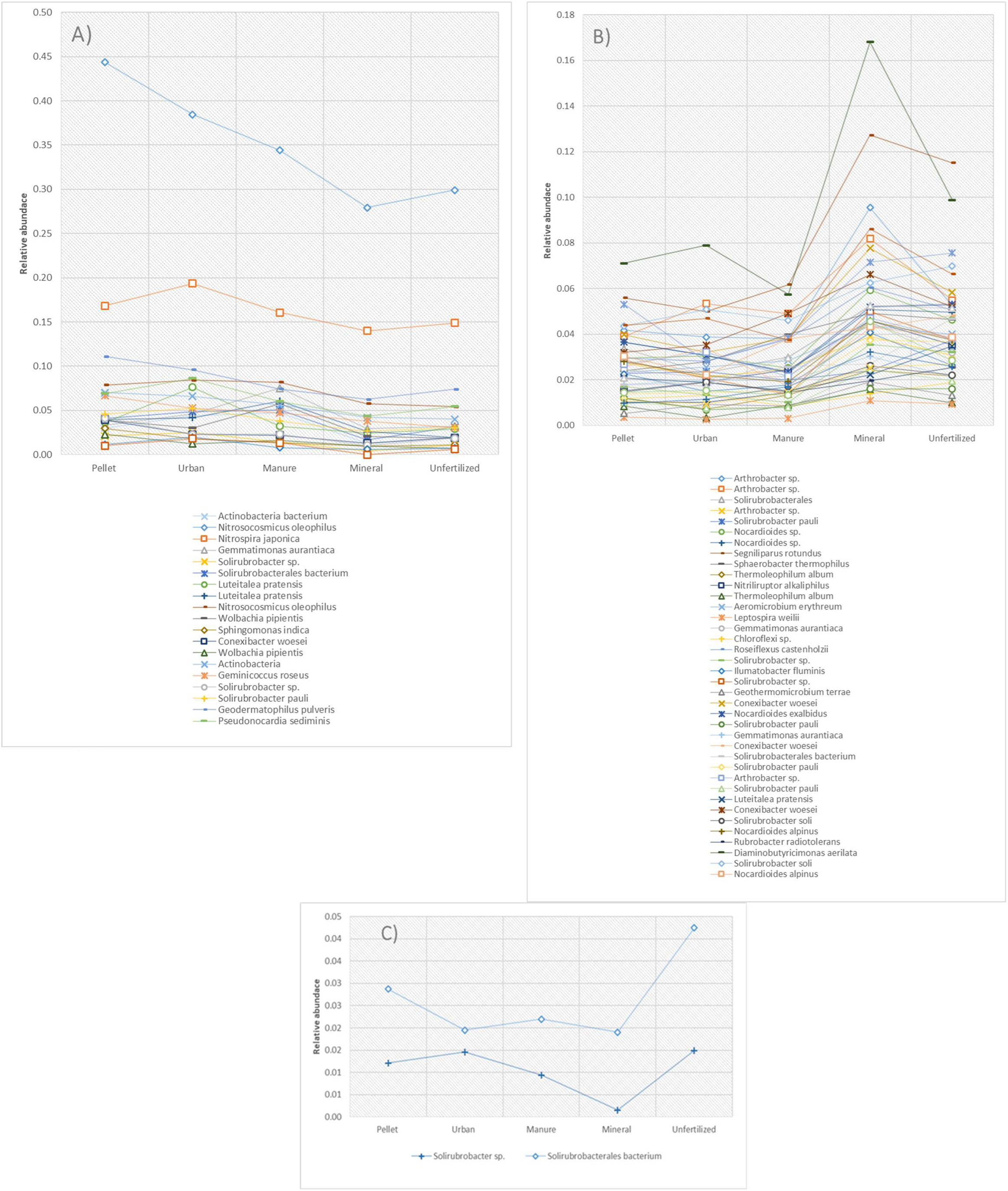
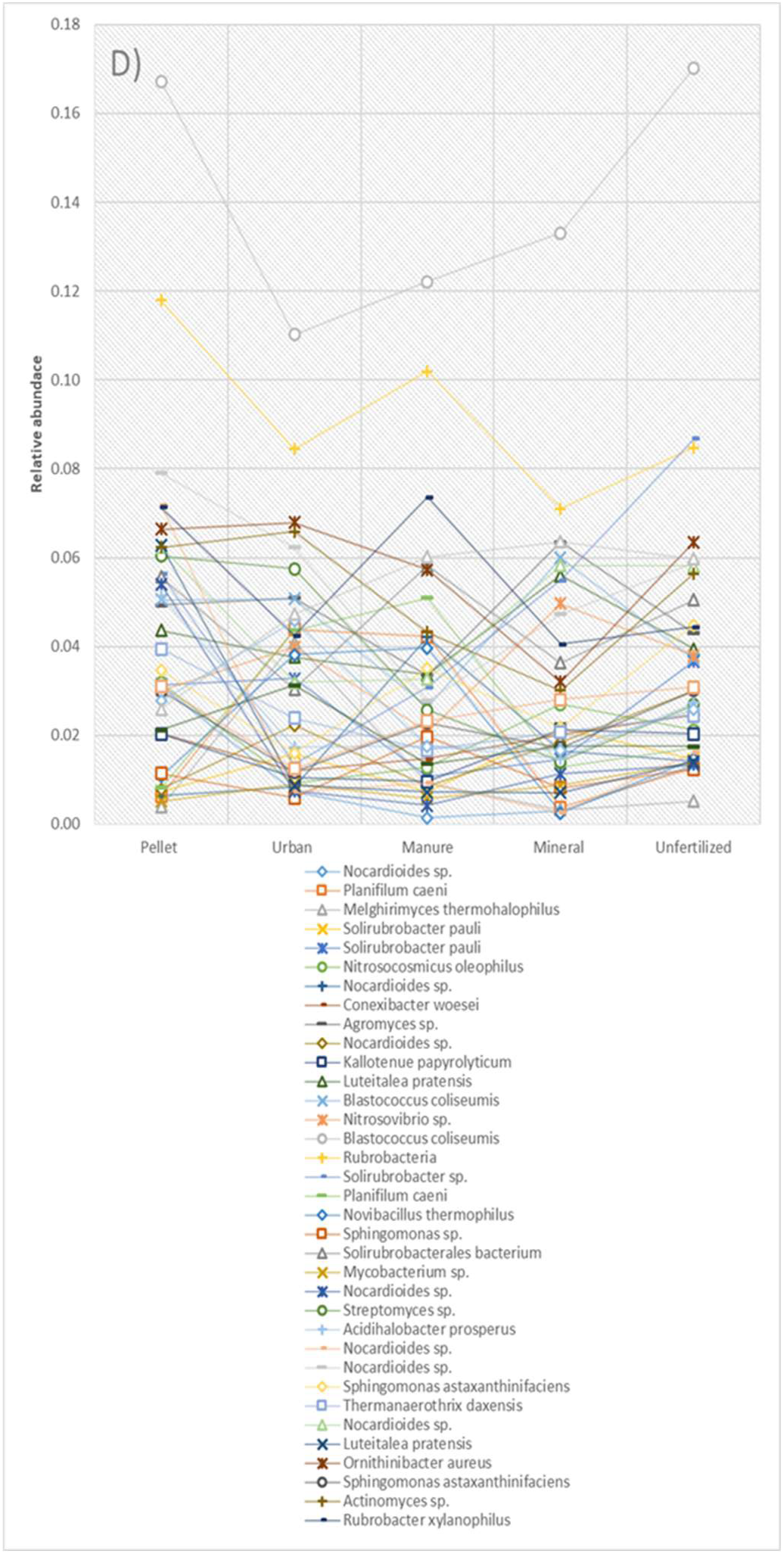
Species groups (SG) showing significantly different abundance values among treatments (average values are plotted). Plot (A) includes 21 SGs with higher average abundance in organic treatments compared to Mineral and Unfertilized treatments. Plot (B) includes 49 SGs with lower abundance in organic treatments compared to Mineral and Unfertilized treatments. Plot (C) includes 2 SGs whose abundance in organic treatments is higher than in Mineral treatment but lower than in Unfertilized treatment. Plot (D) lists 39 SGs where the abundances of the organic treatments differ from each other when compared to Mineral and Unfertilized treatments.

**Supplementary Figure 3.**
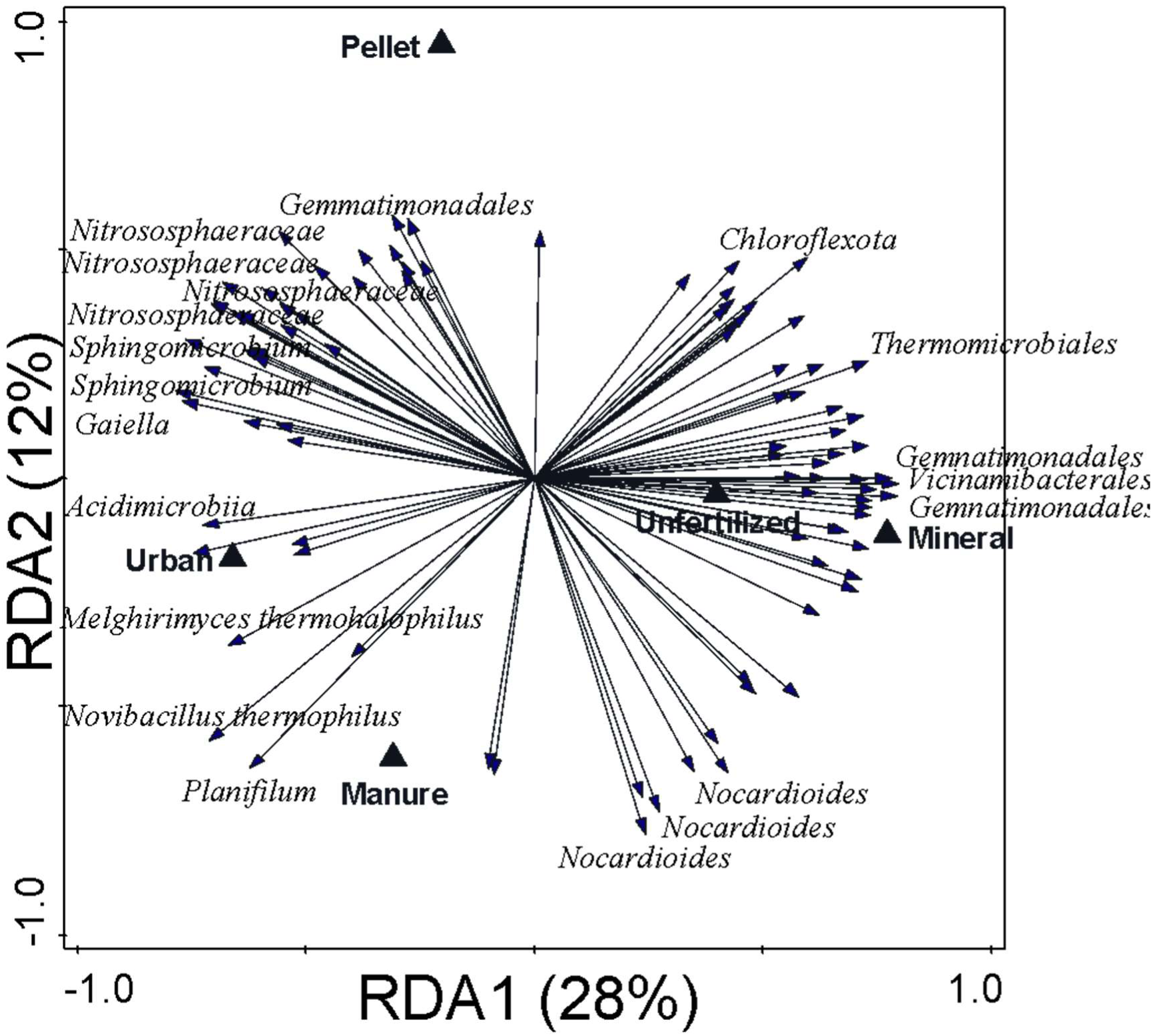
Biplot of the redundancy analysis (RDA) performed on metagenome assembled genome data as response variable and applied treatments as explanatory variables. Only the best-fitting 20 taxa have been labelled. Variance explained by each axis is shown between brackets.

**Supplementary Figure 4.**
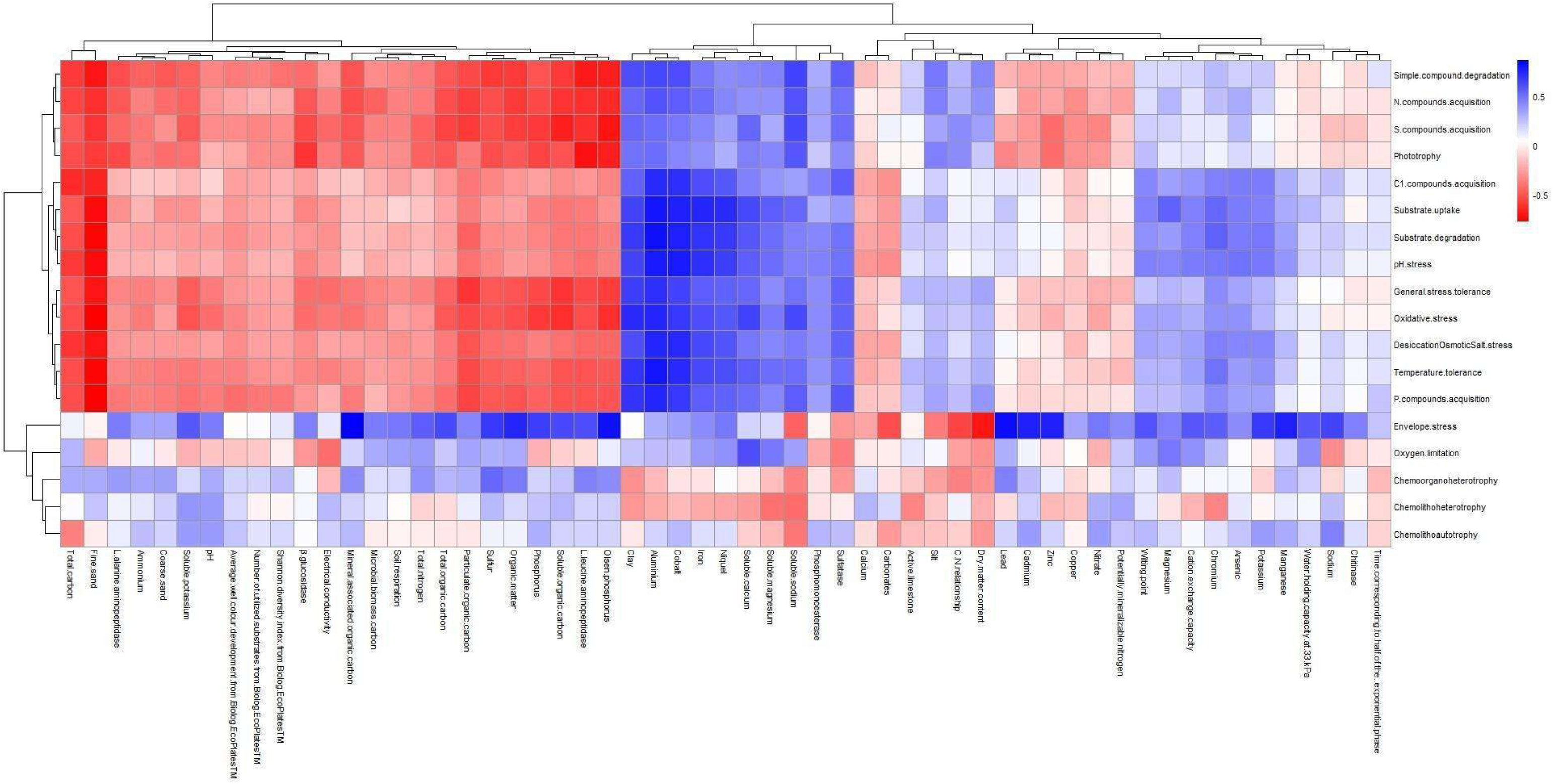
Heat map showing the Spearman correlations between traits at second level of hierarchy and all measured soil properties.

## Notes

### Competing Interest Statement

The authors have declared no competing interest.

